# Co-Electrospinning Extracellular Matrix with Polycaprolactone Enables a Modular Approach to Balance Bioactivity and Mechanics of a Multifunctional Bone Wrap

**DOI:** 10.1101/2025.05.13.653533

**Authors:** Sarah Jones, Madeline Laude, Zeenat Oyebanji, Ruchi Birur, Andres Luengo Martinez, Isabelle Gilbert, Phillip Baek, Elizabeth Cosgriff-Hernandez

## Abstract

Decellularized tissue possesses significant regenerative potential, yet fabricating complex extracellular matrix (ECM) scaffolds remains challenging. Blending with synthetic polymers can aid ECM fabrication but often relies on digested ECM and encapsulation within the synthetic matrix can limit cell-ECM interactions. We recently developed a suspension electrospinning platform to facilitate ECM scaffold fabrication without the need for digestion or polymer-carriers. Its integration into a co-electrospinning system enables modular design of composite scaffolds to combine the regenerative potential of ECM with the advantages of synthetic polymers. This study directly compares co-electrospinning and blend electrospinning of polycaprolactone and small intestinal submucosa (SIS) for use as a bone wrap to augment membrane durability, sustain infection control, and enhance vascularity in Masquelet’s induced membrane technique. Co-spun wraps demonstrated improved handling properties as compared to ECM wraps and solvent welding was used to achieve target suture retention standards without diminishing SIS content. Unlike the blended wraps, the co-spun wraps supported full-thickness cell infiltration within 4 weeks, released gentamicin at a bactericidal concentration for 6 weeks, and demonstrated enhanced angiogenic properties. Collectively, these findings highlight the functionality of a co-electrospinning modular design and demonstrate the efficacy of using a co-spun wrap in bone tissue engineering applications.

## 1. Introduction

Decellularized tissue scaffolds offer a complex make-up of biological cues to support tissue regeneration.^1^ The combination of proteins, proteoglycans, angiogenic growth factors, and matrix-bound vesicles in the extracellular matrix (ECM) guide host cell behavior toward regenerative phenotypes with tissue-specific differentiation and injury resolution.^1-4^ However, advanced manufacturing of ECM materials remains a hurdle with current clinical use of ECM scaffolds limited to the native tissue form or ground powders.^4, 5^ Typically, this limits ECM scaffolds to thin, sheet-type tissues, like the intestine, amnion, and dermis. Current research strategies to develop new form factors include hydrogel development and polymer blending. Hydrogel molding provides geometric flexibility but limited mechanical property ranges that constrain use to non-load-bearing applications. Hydrogel approaches rely on enzymatic digestion to solubilize the ECM into a precursor solution that can denature proteins and limit bioactivity.^6, 7^ Blending ECM with synthetic polymers has enabled more advanced fabrication of ECM-based scaffolds in the form of films, foams, and fibrous materials.^8-12^ Although polymer-blended ECM scaffolds exhibit a broader range of mechanical properties, these handling properties can be compromised with increasing ECM incorporation.^10^ In addition, scaffold blends encapsulate ECM in the secondary polymer matrix which alters tissue remodeling and restricts cell interaction.^11, 12^ There remains a need to fabricate polymer-reinforced ECM-based scaffolds without restricting host interaction with the ECM or compromising handling properties.

Our lab has recently developed a platform for manufacturing fibrous ECM-based scaffolds using suspension electrospinning that does not require polymer carriers. To enable electrospinning, homogenization was used to introduce particle entanglement and elasticity to an ECM suspension. This platform provides a highly tunable framework for designing diverse scaffolds within the broad capabilities of electrospinning. Co-electrospinning is an advanced manufacturing technique that produces scaffolds composed of multiple materials where individual fiber populations are integrated into a single scaffold. Incorporating our ECM suspension into a co-electrospinning setup enables a modular design of a synthetic polymer-ECM composite that maintains the regenerative potential of the ECM fibers and the mechanics of the synthetic fibers. In blended polymer-ECM scaffolds, increasing ECM content raises solution conductivity with a corollary reduction in electrospun fiber diameter, making it difficult to decouple composition from scaffold architectural effects. By contrast, co-electrospinning allows for the decoupling of ECM fiber-mediated cell interactions from the mechanical support provided by the synthetic polymeric fibers.

One potential application of co-electrospinning polymer-ECM composites is the development of multifunctional tissue engineering scaffolds. We previously designed an electrospun polycaprolactone (PCL) wrap to enhance durability and infection control of a bone wrap for Masquelet’s Induced Membrane Technique (MIMT). This bone wrap structurally reinforced the induced membrane, improving suture retention strength by fourfold, and provided antibiotic release for up to eight weeks. However, it lacked a biological component to support tissue regeneration. We hypothesized that integrating ECM fibers into the PCL wrap could guide cell behavior and promote enhanced vascularization of the induced membrane with a corollary benefit on bone regeneration. Decellularized small intestine submucosa (SIS) was selected based on its established role in enhancing vascularization through growth factor release, chemoattractive degradation products, and pro-regenerative macrophage stimulation.^1, 4, 5, 13, 14^ However, incorporating SIS by blending it with PCL in a single electrospinning solution could compromise the diffusion-based release kinetics of gentamicin due to the introduction of hydrophilic components, as well as reduce cell infiltration into the PCL wrap due to reduced fiber diameter. To overcome these challenges, we investigated co-electrospinning of the SIS with PCL into one multifunctional wrap. This approach has the potential to optimize antibiotic release kinetics, handling properties, and angiogenic potential simultaneously. We hypothesized that the SIS fibers will remain accessible for cell interaction and remodeling in the pores between PCL fibers, while the individual PCL fibers will retain their hydrophobicity and slow degradation rate for sustained antibiotic release and mechanical reinforcement.

Herein, we report on the development and testing of a multifunctional bone wrap made of PCL and SIS designed to structurally reinforce the induced membrane, sustain infection control, and enhance vascularity. A co-electrospinning setup was used to investigate the effect of SIS incorporation (10 to 50% by mass) on *in vitro* gentamicin release kinetics, mechanical properties, cell infiltration, and vascularization. A multi-needle system was utilized to increase SIS collection rate to match PCL fiber deposition rates. Co-spun wraps were compared to blended wraps in each study to assess the impact of ECM localization within the wraps. First, the effect of the PCL:SIS compositional ratio on tensile properties was investigated to ensure adequate handling capacity to withstand surgery. The cell infiltration capacity of the wraps was assessed to determine whether membrane integration, necessary for structural reinforcement, was affected by SIS incorporation. Regarding infection control, the antibiotic release kinetics and bacterial inhibition of the composites were evaluated over eight weeks. Finally, the effects of wrap composition on angiogenesis was assessed using a chorioallantoic membrane (CAM) assay. Collectively, these studies highlight the differences between co-spun and blended fiber wraps and demonstrate the potential of the multifunctional wrap to overcome the current limitations of the MIMT.

## 2. Materials and Methods

### 2.1 Materials

All materials were purchased from Sigma Aldrich unless otherwise noted.

### 2.2 Tissue Preparation and Decellularization

The submucosal layer of a 6-9 month porcine small intestine (Animal Technologies) was isolated as described in Ji et al.^15^ The small intestine was first washed in 1X phosphate-buffered saline (PBS) and cut into 10 cm sections. The sections were sliced longitudinally into a flat sheet. The outer layer comprised of the serosa and muscularis externa was removed via mechanical delamination, and the inner mucosa layer was removed via blunted mechanical shearing. The isolated SIS was decellularized with the protocol established by Abraham et al.^16^ First, 10 cm SIS sections were submerged in 100 mM ethylenediaminetetraacetic acid (EDTA) in 10 mM sodium hydroxide (NaOH) for 16 h. Second, the SIS was transferred to 1M hydrochloric acid (HCl) in 1M sodium chloride (NaCl) for 8 h. Third, the SIS was transferred to 1M NaCl in PBS for 16 h. Fourth, the SIS was transferred to PBS for 2 h. Lastly, the SIS was transferred to DI water for 2 h. For each treatment, the 10 cm SIS sections were submerged in 5 mL of solution each and continuously agitated at 37ºC. The prepared SIS was stored at -80ºC until further use.

### 2.3 Fabrication of Co-spun Wraps

A co-electrospinning setup was used to fabricate fibrous wraps with isotropic orientation composed of PCL and SIS as depicted in **Figure 1**. The gentamicin-loaded PCL spinning solution was prepared from PCL with a molecular weight of 80,000 g/mol. First, gentamicin was dissolved at a concentration of 11 mg/mL in hexafluoroisopropanol (HFIP) with continuous agitation at 37ºC for 2 weeks. Subsequently, PCL was dissolved in the gentamicin solution at 12 wt% of HFIP for resulting dry fibers of 5 wt% gentamicin. SIS electrospinning suspensions were prepared as described in Jones et al.^17^ Briefly, the SIS-derived ECM was submerged in DI water and thawed at 37 ºC. The SIS sheets were then homogenized with an OMNI Tissue Homogenizer TH (OMNI International) for 20 - 30 s. The homogenized SIS was flash-frozen in liquid nitrogen and lyophilized for 2 days. The dried SIS was broken into particles with an electric blade grinder (Kaffe). SIS particles < 2000 µm in diameter were suspended in chilled HFIP at 40 mg/mL and allowed to mix for over 12 h at 4 ºC. Immediately prior to electrospinning, the SIS suspensions were homogenized for 20 - 30 s.

**Figure 1:**
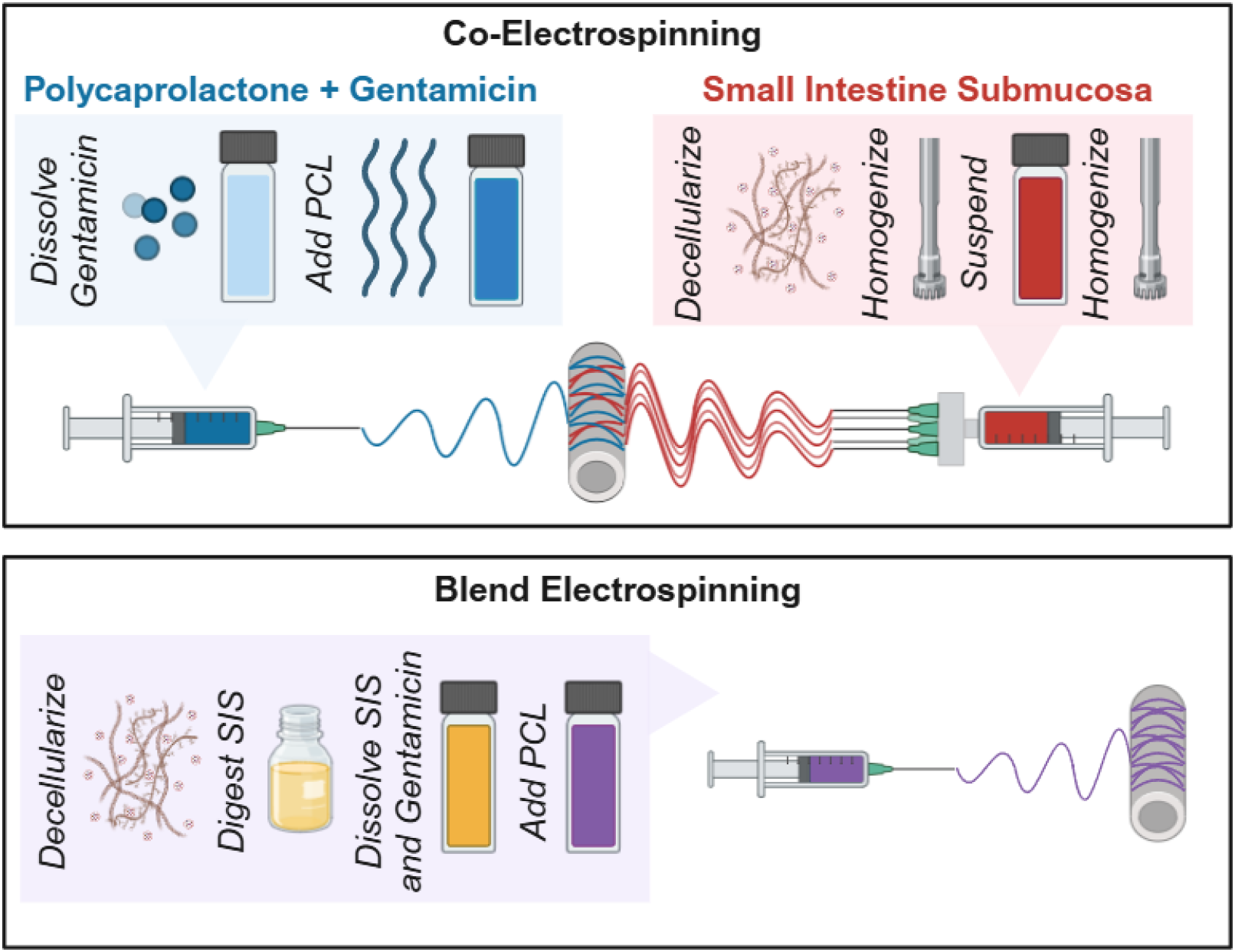
Schematic diagram comparing the protocols for co-electrospinning and blend electrospinning of PCL:SIS wraps. Created in BioRender. Jones, S. (2025) https://BioRender.com/kx4epno

For the PCL control wraps, the PCL solution was pumped from a 20 G blunted needle charged with 8 kV at a flow rate of 1 mL/h. The needle was set up 12 cm from a grounded cylindrical collector covered in parchment paper rotating at 50 rpm. For the SIS control wraps, the SIS suspension was pumped from a single 20 G blunted needle charged with 18 kV at a flow rate of 0.5 mL/h. The needle was set up 20 cm from a grounded copper plate covered in parchment paper. All wraps were vacuum- dried to remove excess solvent before further testing. For the co-spun PCL:SIS wraps, the PCL solution was loaded into a single syringe positioned 12 cm from the collector. The SIS suspension was divided equally into 5 syringes positioned on a single syringe pump placed 15 cm from the collector, directly opposite from the PCL syringe placement. The PCL solution was pumped from a 20 G blunted needle charged with 10-13 kV at a flow rate of 1 mL/h. Simultaneously, the SIS suspension was pumped from the 18 G blunted needles charged with 19 kV at a varying flow rate dependent on the desired PCL to SIS collection ratio. For 90:10, 70:30, and 50:50 PCL:SIS wraps, a SIS flow rate of 0.5, 1.5, and 3 mL/h was used. Each wrap was electrospun on a cylindrical mandrel covered in parchment paper rotating at 50 rpm and charged with - 4 kV. Relative humidity was maintained between 30-50%.

### 2.4 Fabrication of Blended Wraps

For blended wraps, the decellularized SIS was first lyophilized and ground into a powder with an electric blade grinder (Kaffe) and then digested in pepsin to facilitate solubility in HFIP. Briefly, the SIS powder was submerged at a concentration of 10 mg/mL in 3% acetic acid with 1 mg/mL pepsin and stirred at 37 °C for 72 hours. The solution was neutralized with 10 M NaOH and passed through a 150- mesh filter. The solution was diluted 2-fold and lyophilized. Blended PCL:SIS solutions were prepared as a 10 wt% solution in HFIP with a 90:10, 70:30, and 50:50 mass ratio of PCL:SIS. Gentamicin was loaded at a concentration of 5 wt% of PCL. First, gentamicin was dissolved at 11 mg/mL in HFIP at 37 °C for 2 weeks. The gentamicin solution was diluted to the desired concentration, and SIS was dissolved in the gentamicin solution overnight at 4 °C. The SIS solution was passed through a 400-mesh filter to remove non-dissolved particles. Lastly, PCL was dissolved into the SIS:gentamicin solution overnight at 4 °C.

To electrospin the blended wraps, the PCL:SIS solution was loaded into a single syringe positioned 20 cm from the collector. The solution was pumped from a 20 G blunted needle at a flow rate of 1 mL/h. For 90:10, 70:30, and 50:50 PCL:SIS wraps, the needle was charged at 13 kV, 13 kV, and 18 kV, respectively. Each wrap was electrospun on a cylindrical mandrel covered in parchment paper rotating at 50 rpm and grounded. Relative humidity was maintained between 30-50%.

### 2.5 Wrap Composition Characterization

To characterize fiber morphology, wrap specimens (n = 3) were coated with a 5 nm layer of gold (Sputter Coater 108, Cressington Scientific Instruments) and imaged with scanning electron microscopy (SEM, Phenom Pro, NanoScience Instruments, Phoenix, AZ) at an accelerating voltage of 10 kV. Five micrographs were taken of each wrap in a raster pattern (n = 3). The fiber diameter distribution was determined by measuring 100 fibers in each mesh with ImageJ® Software.

Changes in wrap composition were characterized by attenuated total reflectance-Fourier transform infrared (ATR-FTIR) spectroscopy. Spectra were collected on a Thermo Scientific Nicolet iS10 FTIR equipped with a germanium crystal at a resolution of 2 cm^-1^ for 32 scans. Five spectra were collected from each wrap in a raster pattern (n = 5). PCL content within co-spun wraps was characterized using relative peak height analysis by normalizing the 1724 cm^-1^ (C=O ester) peak height in co-spun or blended wraps to the corresponding peak from the PCL control wrap. Similarly, SIS content within co-spun and blended wraps was characterized by normalizing the 1648 cm^-1^ peak (C=O amide I) to the corresponding peak from the SIS control wrap.

### 2.6 Mechanical Assessment and Solvent Welding

To assess the handling properties of the wraps, uniaxial tensile and suture retention testing were conducted. For tensile testing, a die was used to cut electrospun wraps into microtensile specimens with a gauge length of 13 mm and gauge width of 5 mm in accordance with the ASTM method D1708. Dogbone specimens were strained at a rate of 100%/min based on the initial gauge length until failure (Instron 2712-019). Stress-strain plots were generated to determine the tensile properties of the wraps including 2% secant modulus, ultimate elongation, and tensile strength. To evaluate suture retention strength, the wraps were cut into 10 × 20 mm rectangles. A 2/0 Nylon suture was placed with a 2 mm bite centered along the major axis, and the suture was stretched at a rate of 50 mm/min until failure (Instron 2712-019). The max force exhibited during suture pull-out testing was identified as the suture retention strength of the sample. Studies were conducted with n = 9 of co-spun or blended wraps. All mechanical assessments were conducted on dry specimens at room temperature. The SIS control wrap (0:100 PCL:SIS) could not be measured with confidence, so the data was not reported. The 10 N load cell utilized for the composite wraps was not sensitive enough to precisely measure the suture retention strength.

Solvent welding was utilized to selectively increase PCL fiber fusion within the co-spun wraps to increase the suture retention strength. Briefly, a basin filled with 50 mL of chloroform was placed in a glass desiccator and covered with a metal mesh (0.5 cm^2^ pores). The 70:30 and 50:50 PCL:SIS co- spun wraps were placed on the metal mesh and sealed in the desiccator for 13 - 15 min, the maximum time before bulk mesh changes was observed. The wraps were subsequently dried on the benchtop for at least 8 hours. The chloroform exposure and drying sequence was repeated 3 times. The samples were vacuum dried overnight before undergoing mechanical testing.

### 2.7 Cell Attachment and Infiltration

*In vitro* cell attachment and infiltration were assessed as an initial measure of the capacity of each wrap to support tissue integration. The wraps were first vacuum-dried to remove any residual HFIP after electrospinning. The wraps were cut into 8 mm discs and sterilized in 70% ethanol for 3 hours. A wetting ladder was conducted by incremental changes of 50%, 30%, and 0% ethanol in PBS for 30 min each, and then soaked in PBS overnight before cell seeding. Human dermal fibroblasts (hDFs) were expanded in α-MEM growth media, supplemented with 10 % fetal bovine serum (Atlanta Biologicals, Flowery Branch, GA) and 1% of penicillin-streptomycin (10,000 U/mL, Thermo Fisher Scientific) at 37 ºC and 5 % CO^2^. To assess cell attachment, fibroblasts were seeded on the surface of the wraps at 10,000 cells/cm^2^. After incubation for 3 hours, the cells were fixed in 3.4 % glutaraldehyde. The nuclei on the surface were stained with SYBR green (Biotium). Five fluorescent images were taken of each wrap in a raster pattern (n = 4). The number of cells per image were counted with respect to area. To assess cell infiltration, the fibroblasts were seeded on the surface of the wraps at 100,000 cells/cm^2^. The samples were cultured for 4 weeks and fixed in 3.4 % glutaraldehyde at weekly timepoints. The wraps were embedded in 5% w/w gelatin and 5% w/w sucrose and cryo-sectioned axially with a 10 μm thickness to analyze a cross- section of the wraps. The nuclei throughout the cross-section were stained with SYBR green (Biotium) and fluorescently imaged. Four cross-sections were imaged per sample (n = 4). Pixels within ± 25 from the RGB value of 19, 207, 27 were identified as nuclei and the distance from the seeding edge of the wrap to each pixel was measured with a custom code provided in the data repository. A histogram of cell distribution was compiled from these measurements. The maximum infiltration depth that 90% of the cells was distributed was also calculated. Studies were conducted with n = 4 replicates.

### 2.8 Gentamicin Release Profile

The release kinetics of gentamicin of the co-spun and blended wraps were compared to the PCL control wrap. Briefly, specimens were submerged in 1 mL of DI water to simulate sink conditions and incubated with agitation at 37 °C for 8 weeks. After 24 hours and weekly timepoints, the release medium was collected and replaced with fresh DI water. Releasates were stored at - 4 °C until bacteria culture was conducted. The residual proteins from SIS in the releasate prohibited the use of standard Ninhydrin assay to quantify gentamicin release. Gentamicin concentration in the releasate was analyzed with an agar diffusion assay as described previously.^18^ Briefly, osteomyelitis-specific methicillin-susceptible Staphylococcus aureus (ATCC®, strain 49230) was suspended in brain heart infusion (BHI) broth and incubated while mixing at 240 rpm and 37 ºC overnight. The inoculate was diluted to 2.0 × 10^8^ CFU/mL to match the turbidity of a 0.5 MacFarland standard.^19^ The bacterial suspension was spread evenly onto BHI agar, then wells with a 6 mm diameter were punched from the agar. The wells were filled with 50 μL of each release sample. The plates were incubated at 37 ºC overnight and the diameters of the zones of inhibition were measured with ImageJ. The concentration of gentamicin released at each timepoint was then calculated using a standard agar diffusion curve of gentamicin in DI water with a concentration range of 0 – 800 μM. Three bacteria culture replicates were conducted for each wrap (n = 3).

### 2.9 Zone of Bacterial Inhibition

The bactericidal activity of the released gentamicin concentration was analyzed with the Kirby-Bauer zone of inhibition assay.^19^ Osteomyelitis-specific methicillin-susceptible Staphylococcus aureus (ATCC®, strain 49230) was suspended in brain heart infusion (BHI) broth and incubated with agitation at 37 ºC overnight. The inoculate was diluted to 2.0 × 10^8^ CFU/mL to match the turbidity of a 0.5 MacFarland standard.^19^ The bacterial suspension was spread evenly onto BHI agar, and samples cut to 6 mm discs were immediately placed on the surface. The agar plates were incubated while stationary at 37 ºC for 16 – 18 h. The diameters of the zones of inhibition were then measured. Studies were conducted with an n = 4.

### 2.10 In Ovo Angiogenesis

A chorioallantoic membrane (CAM) assay was used to measure the angiogenic capacity of the wraps. Co-spun and blended PCL:SIS wraps were compared to a 200 µm Nylon mesh negative control. Briefly, fertilized Japanese quail eggs (Bryant’s Roost) were incubated horizontally at 37 ºC and 50% relative humidity for 3 days. The eggshells were sterilized with 70% ethanol and allowed to dry. The embryo was removed from the shell and placed in a 50 mm petri dish. Three 50 mm polystyrene dishes were placed in a 150 mm petri dish filled with 5 mL DI water to maintain a 100% humid environment and incubated at 37 ºC for 7 days. The samples were cut to 6 mm discs and placed 2 – 3 mm to the side of the central anterior vitelline vein in the outer third of the CAM as previously described.^20^ Images were taken daily for 4 days with a stereoscope to visualize angiogenesis. Vessel density was quantified as the change in the number of vessels crossing the outer perimeter of the sample from Day 0 to Day 4 (n = 3 - 6).

### 2.11 Statistical Analysis

Data are represented as mean ± standard deviation unless stated otherwise. Statistical Analysis was performed utilizing ordinary one-way ANOVA with Tukey’s post hoc analysis in GraphPad Prism 10 unless stated otherwise. Statistical significance was accepted at p < 0.05; no marking indicates p > 0.05; * indicates p < 0.05; ** indicates p < 0.01; *** indicates p < 0.001; **** indicates p < 0.0001. All groups were compared to the 100:0 PCL:SIS control unless stated otherwise. Outliers were removed using Grubbs’ analysis (alpha = 0.01).

## 3. Results and Discussion

Building upon our prior work fabricating a durable and antimicrobial PCL bone wrap for use in MIMT,^21^ we utilized our new suspension electrospinning method to incorporate SIS into the wrap to enhance vascularization. A co-electrospinning setup was designed to generate a multifunctional wrap with individual PCL and SIS fiber populations. The differing collection rates of PCL and SIS posed a challenge to incorporating a substantial volume of SIS. When electrospun alone, a PCL spinning solution has a concentration over 200 mg/mL and is pumped from a syringe pump at 1 mL/h; whereas, the SIS spinning suspension has a concentration of 40 mg/mL and is pumped at 0.5 mg/mL. When each material was electrospun in parallel onto a single mandrel with its original spinning conditions, the wrap was over 95% PCL. Significantly changing the flow rate or concentration of either material risked sacrificing the morphology and spinnability of the PCL and SIS, respectively. To address this issue, a multi-jet system was designed to electrospin five SIS jets simultaneously with one PCL jet, **Figure 1**. A custom mount for 5 syringes was 3D printed to stabilize the SIS syringes on a single syringe pump positioned on one side of the collecting mandrel. A positively charged wire was wrapped around each needle to apply a consistent charge. The single PCL syringe was set up opposite of the collecting mandrel from the SIS syringes. This setup facilitated a comparable collection rate of PCL and SIS, and could be easily modified to achieve a range of PCL:SIS compositional ratios. Initially, 90:10, 70:30, and 50:50 PCL:SIS wraps were electrospun with a 0.5, 1.5, and 3 mL/h flow rate from the 5 SIS needles. However, the SIS fibers in the 90:10 wrap were much smaller than in the 70:30 and 50:50 wraps, so an alternative method of using 1 SIS syringe with 1.5 mL/h flow rate was adopted to achieve more consistent fiber morphology across the groups, **Figure 2A**. The resulting wraps exhibited fibers with 3 fiber diameter clusters around 0.2 µm, 1 µm, and 5 µm. The increasing presence of smaller 0.2 and 1 µm fiber populations corresponded with increasing SIS incorporation and matched the morphology of the fibers in the SIS wrap. Similarly, the 5 µm fiber population matched the morphology of the fibers in the PCL wrap. The complete fiber diameter distribution of the wraps is provided in **Figure S1**.

**Figure 2:**
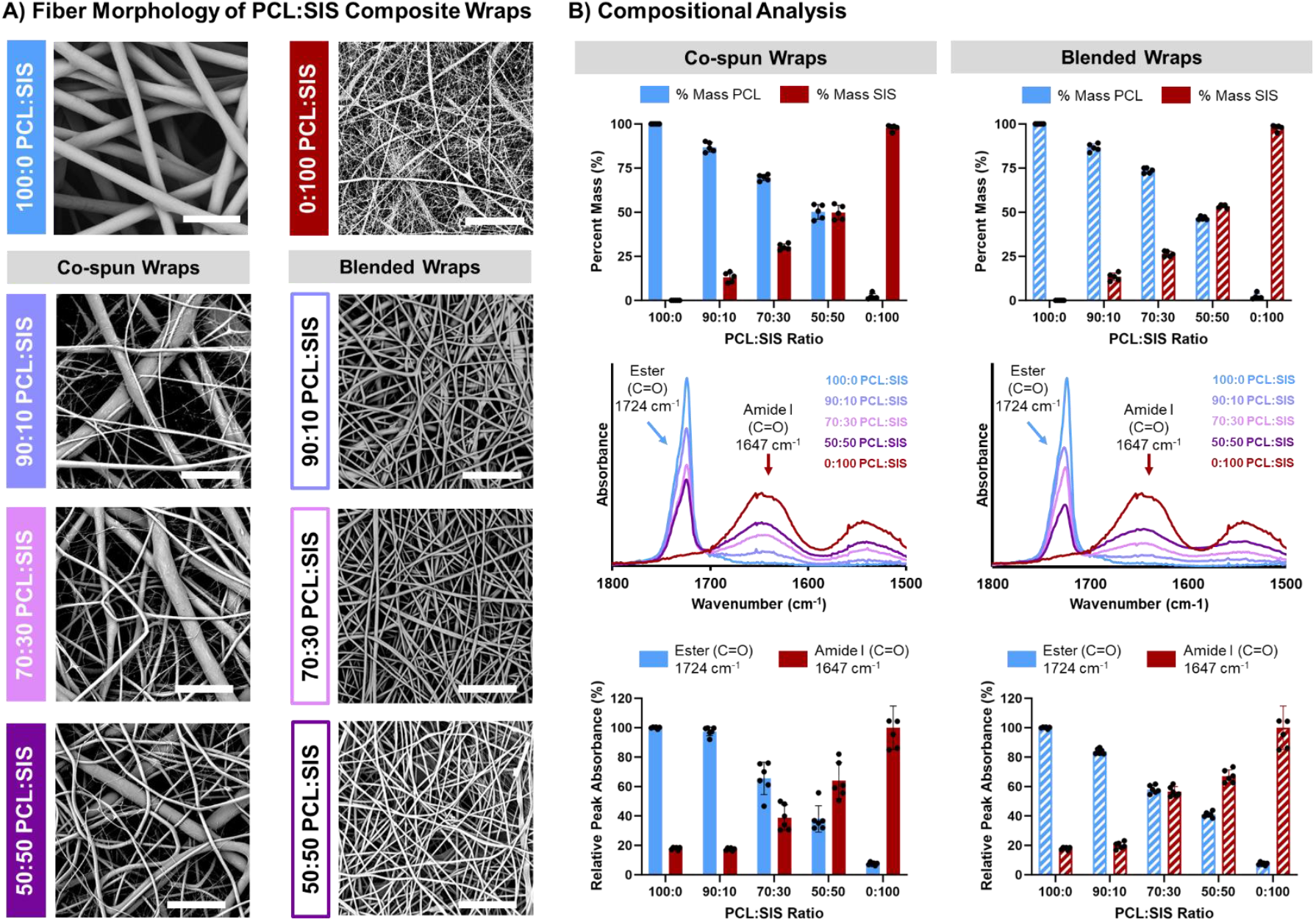
Co-spun and blended PCL:SIS wraps fabricated with target compositional ratios 100:0, 90:10, 70:30, 50:50, and 0:100 PCL:SIS. A) Scanning electron microscopic images of fiber morphology. Scale bar = 20 µm. B) Compositional analysis of co-spun and blended wraps including mass fraction in each wrap from selective dissolution and attenuated total reflectance-Fourier transform infrared spectral (ATR-FTIR) analysis normalized to control PCL and SIS peak heights. Full spectra are available in the supplemental information.

Blended wraps with matching PCL:SIS compositional ratios were electrospun following methods established in the literature, **Figure 1**.^22, 23^ To electrospin a blended wrap with over 10% SIS, the SIS must first be digested to facilitate dissolution in the electrospinning solution. Attempts to electrospin a PCL solution loaded with non-digested fine SIS particles resulted in non-uniform and split fibers with minimal SIS incorporation in the resulting electrospun meshes. Digestion in pepsin facilitated SIS solubility in the blended electrospinning solution and resulted in a more homogenous fiber distribution. The 10 wt% solution made from digested SIS and PCL resulted in a wrap with a fiber diameter of 0.8 ± 0.1 µm, similar to the SIS fiber diameter in the co-spun wraps. Efforts to increase the fiber diameter near the PCL fibers in the co-spun wrap were unsuccessful. Increased concentrations up to 15 wt% resulted in wraps with a wide fiber diameter distribution (1.9 ± 1.2 µm), still under 2 µm. The high conductivity of the added SIS likely introduced instability in the electrospinning jet that was unable to withstand the lower surface tension needed to electrospin larger fibers without splitting. The blended electrospinning troubleshooting results are detailed in **Figure S2**. To our knowledge, there are no reports of electrospun polymer:ECM blended wraps with a fiber diameter greater than 2 µm, which is further limited to below 1 µm with over 10% ECM incorporation.^10, 22-28^ Thus, to maintain a more uniform fiber diameter distribution that is characteristic of blend electrospun meshes in literature, the blend spinning solutions were made by dissolving PCL, gentamicin, and pepsin-digested SIS in HFIP at a 10 wt% concentration. PCL and SIS were mixed at mass ratios that correspond to the 90:10, 70:30, and 50:50 PCL:SIS co-spun wrap compositions. The blended wraps exhibited a more uniform fiber morphology compared to the co-spun wraps, **Figure 2A**. Additionally, the diameter of the blended fibers decreased from 1.3 ± 0.4 to 0.6 ± 0.3 µm with increasing SIS incorporation from 10 to 50% due to reduced viscosity and increased conductivity of the spinning solutions, **Figure S1**.

A selective mass loss experiment was designed to quantify the relative mass ratio of PCL and SIS in both co-spun and blended wraps, **Figure 2B**. Toluene was used to selectively dissolve the PCL from the co-spun and blended wraps. The dry mass of the wraps before and after submersion in toluene was used to calculate the mass fraction of PCL and SIS. As expected, the co-spun wraps that were electrospun with an increased SIS flow rate and more SIS syringes exhibited an increase in SIS mass fraction and reduced PCL mass fraction. Similarly, the blended wraps that were electrospun from solutions with increased SIS concentration exhibited an increased SIS mass fraction and reduced PCL mass fraction in good agreement with target ranges. ATR-FTIR was also performed to validate the chemical composition of the composite wraps, **Figure 2B**. Full ATR-FTIR spectra from 800 – 4000 cm^- 1^ are shown in **Figure S3**. The characteristic ester peak corresponding to C=O stretch present at 1724 cm^-1^ was used to estimate relative concentration of PCL. The characteristic amide I peak corresponding to the C=O stretch at 1647 was used to estimate the relative concentration of SIS. In both the co-spun and blended wraps, the height of the ester peak decreased and the height of the amide I peak increased with increasing SIS incorporation. Relative peak height analysis with normalization to control PCL and SIS peak heights indicated compositional ranges in good agreement with mass fractions and target ranges. Collectively, these results indicate that the multijet co-electrospinning setup was successful in achieving a broad range of PCL:SIS compositions (90:10, 70:30, and 50:50 PCL:SIS) and enabled a rigorous analysis of the impact of SIS concentration and localization on scaffold performance.

### 3.1 Effect of SIS Incorporation on Wrap Mechanics

The impact of composition on wrap mechanical properties was evaluated using a suture retention test and uniaxial tensile test, **Figure 3**. Full stress-strain plots are available in **Figure S4**. Co- spinning the SIS with PCL increased its tensile properties and suture retention strength. The SIS wrap control was relatively brittle with low ultimate elongation and suture retention strength that could not be measured. The co-spun PCL:SIS wraps exhibited increased suture retention strength and increased tensile strength and ultimate elongation with increasing PCL incorporation. The 70:30 and 50:50 PCL:SIS co-spun wraps did not meet the ISO 7198 suture retention standard (> 2.0 N), but were 2-fold higher than the induced membrane. Two methods were explored to increase suture retention without limiting SIS incorporation. It was found that increasing the thickness of the 50:50 PCL:SIS wrap from 300 µm to 600 µm increased the suture retention strength of the wrap above 2.0 N (data not shown). However, increasing the thickness of the wrap risks could potentially limit cell infiltration through the full depth of the wrap before bone graft placement in the MIMT. Therefore, solvent welding was explored as a way to increase suture retention strength by selectively increasing PCL fiber fusion post-fabrication without impacting SIS morphology. The 70:30 and 50:50 PCL:SIS co-spun wraps after solvent welding exhibited suture retention strength similar to the PCL wrap and achieving ISO 7198 suture retention standards, **Figure 3C**. These results highlight the high tunability that is facilitated by the modular approach of co-electrospinning. The PCL network could be modified to increase the handling properties of the wrap without impacting SIS incorporation or morphology. In contrast to co-spun wraps, each blended wrap composition met the ISO 7198 suture retention standards but with greater variability than the co-spun wraps. It is expected that the reduced fiber diameter in the blended wraps compared to the PCL wrap and co-spun wraps contributed to the increased suture retention strength due to increased fiber interconnection points and density. Interestingly, the suture retention strength did not follow a linear trend with increasing SIS incorporation, as the 70:30 blended wrap demonstrated the highest strength, **Figure 3B**. Altogether, SIS incorporation increased the brittle behavior of both the co-spun fiber and blended wraps, but all of the wraps exhibited elevated handling properties as compared to the induced membrane and could be designed to meet suture retention standards.

**Figure 3:**
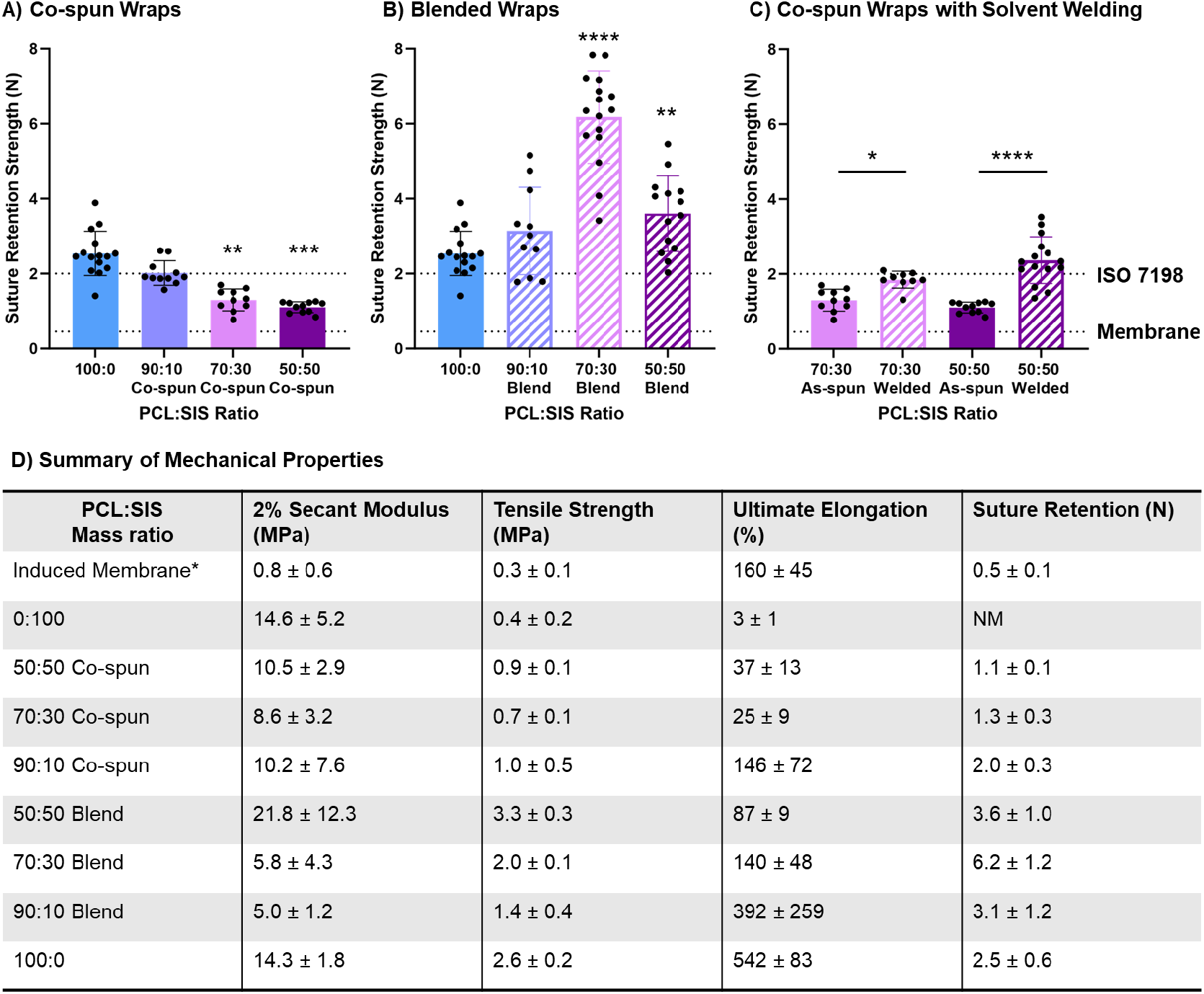
The impact of SIS incorporation on the mechanical properties of co-spun and blended PCL:SIS wraps. Suture retention strength of as-spun co-spun wraps (A) and blended wraps (B) compared to the ISO 7198 standard (> 2.0 N) and the induced membrane (0.46 N). C) Suture retention strength of solvent vapor welded co-spun wraps. A-C) Dotted lines represent the ISO 7198 standard (top) and the induced membrane (bottom). D) The 2% secant modulus, tensile strength, ultimate elongation, and suture retention strength of PCL:SIS wraps. NM = not measurable with instrument. *Reported values for 2% secant modulus, tensile strength, and ultimate elongation are from Gaio et al.^29^

### 3.2 Effect of SIS Incorporation on Cell Infiltration

As an initial assessment of the potential for tissue integration, cell attachment was evaluated by culturing fibroblasts on the surface of the wraps. All substrates demonstrated statistically similar (p > 0.05) levels of cell attachment after 3 hours and were elevated as compared to the tissue culture polystyrene control, **Figure S5**. *In vitro* cell infiltration was then assessed by seeding fibroblasts on the surface of the wraps and cross-sectional analysis of cell migration into the wrap monitored over four weeks, **Figure 4**. All co-spun wraps displayed full cell infiltration over the four weeks with similar or greater cell distribution as the PCL control. Additionally, the total area of cell nuclei within the wraps is greater in the co-spun wraps as compared to the PCL control, which suggests the SIS promotes increased cell proliferation, **Figure 4D**. These results were somewhat unexpected. We have previously shown that PCL wraps fabricated with reduced fiber diameter exhibited reduced cell infiltration capacity.^21^ This was attributed to the reduced pore size estimated from decreased fiber diameter. Given that the average pore size predicted by the bimodal fiber diameters in co-spun wraps indicated a marked decrease in pore size (**Figure 4E**), there was a concern that the presence of SIS fibers that were less than 1 µm in diameter in the co-spun wraps could reduce cell infiltration. However, more fibroblasts infiltrated over 50 µm into the wraps with increasing SIS incorporation, which indicates that reduced pore size caused by the smaller SIS fibers (< 2 µm in diameter) in the co-spun wraps did not impede cell migration, **Figure 4E-F**. These results are opposite from Pham et al. who demonstrated that a PCL electrospun wrap with a mixture of micro and nanofibers inhibited cell infiltration compared to a microfiber control.^30^ This suggests that cell migration is functionally different on SIS and PCL fibers, possibly due to differences in fiber stiffness. We hypothesize that the fibroblasts can navigate around and/or manipulate SIS fibers similar to native ECM structures, and the bioactive cues provided by the SIS components promoted cell migration and proliferation within the wrap.

**Figure 4:**
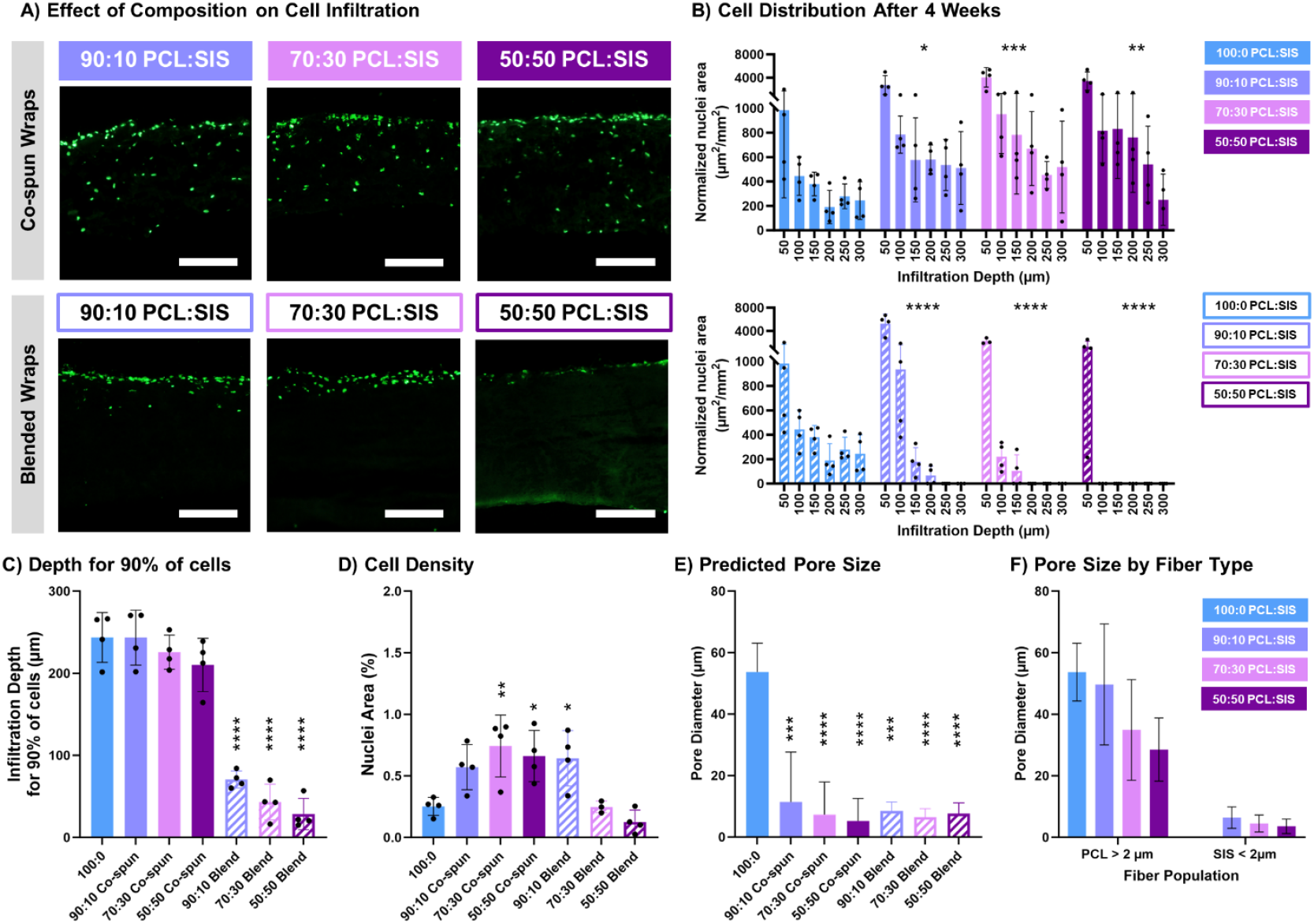
Cell infiltration into PCL:SIS wraps after 4 weeks of culture. A) Cross-sectional images of cell distribution. Scale bar = 200 µm. B) A histogram of cell distribution across the wrap depth after 4 weeks. The x-axis labels are the bin maximum for each column. C) The maximum infiltration depth of 90% of the cells that are distributed throughout the wrap. D) The percent area of nuclei in the wrap cross-section. E) The pore size of the wraps calculated from average fiber diameter and porosity. F) The pore size of the co-spun wraps corresponding to the PCL and SIS fiber populations calculated separately for fibers less than and greater than 2 µm.

In contrast, fibroblasts were largely limited to the surface of the blended wraps. Limited cell infiltration was seen in the 90:10 PCL:SIS wrap, and infiltration depth was reduced with increasing SIS incorporation. The lack of cell migration into the blended wraps was largely attributed to the smaller fiber diameter of the wraps. Although the cells were capable of migrating past the < 1 µm SIS fibers in the co-spun wraps, the cells were unable to navigate through the PCL:SIS blended fibers of a similar diameter. Considering the overall pore size of the co-spun and blended wraps are similar **(Figure 4E)**, this was attributed to the differing composition of the fibers themselves. The pores created by the PCL fibers in the co-spun wraps are at least 3 times larger than the pores between the composite blended fibers, **Figure 4F**. Although the blended wraps did not support significant cell migration, the 90:10 PCL:SIS wrap exhibited significantly elevated nuclei area in the top 50 µm of the wrap compared to the PCL control, indicating that the fibroblasts readily proliferated on its surface. No impact on cell proliferation was observed on the 70:30 and 50:50 PCL:SIS blended wraps. Collectively, co-spun wraps displayed superior cell infiltration capacity compared to the blended wraps and supported cell infiltration through the full thickness of the wrap after 4 weeks at all PCL:SIS ratios.

### 3.3 Effect of SIS Incorporation on Gentamicin Release

The impact of SIS incorporation on infection control of the composite wraps was assessed through gentamicin release kinetics and bactericidal activity. Gentamicin release kinetics is commonly detected with a ninhydrin assay that detects amines.^21, 31^ However, the SIS component in the composite wraps contains amines that interfere with the ninhydrin detection of gentamicin. Thus, an agar diffusion assay was conducted, similar to prior work.^18^ This assay directly measures the bactericidal activity of the release medium compared to a standard curve and is not impacted by the presence of SIS. The agar diffusion assay was used to measure the release of gentamicin from the composite wraps weekly for 8 weeks. Each wrap was loaded with gentamicin at a concentration of 5 wt% of PCL mass. Thus, with decreasing PCL incorporation, less gentamicin was loaded into the wrap, **Figure 5A**. Despite the reduced loading concentration of gentamicin with increased SIS incorporation, the co-spun wraps maintained a released dose above the minimum bactericidal concentration (MBC) for 6 weeks, **Figure 6A**. After 6 weeks, the dose was reduced to below the MBC but was maintained above the minimum inhibitory concentration (MIC) for 8 weeks. The combination of reduced loading and maintained dose release resulted in a higher efficiency of release from the co-spun wraps, where the 50:50 PCL:SIS co- spun wrap released 72 ± 6% of the loaded gentamicin compared to 35 ± 2% from the PCL wrap, **Figure 5B**. This is in stark comparison to 1.5 ± 0.7% from a cement spacer clinical control.^21^ Gentamicin is delivered from the PCL fibers via diffusion. Therefore, the hydrophilicity of the wrap will impact the gentamicin release kinetics. A swelling ratio assessment confirmed that the co-spun fiber and blended wraps with elevated SIS levels exhibited a higher water uptake percentage by mass **(Figure 5C)**. Thus, we hypothesize that the added hydrophilicity instilled by the SIS incorporation overcomes the relative mass loss of PCL and increases gentamicin diffusion from the composite wraps. The blended wraps demonstrated similar release trends at early time points, **Figure 6A**. The 24-hour burst release was elevated with increasing SIS incorporation, and the weekly release was similar to PCL for the first two weeks. Thus, the percent yield of gentamicin release was also elevated with increasing SIS incorporation, with an exception for the 90:10 blended wrap. However, the gentamicin release from the 90:10 and 70:30 blended wraps dropped below the MBC after 3 and 4 weeks, respectively. This was attributed to the reduced fiber diameter and elevated surface area to volume ratio of the blended wraps compared to the PCL wrap, causing reduced release longevity. However, the added hydrophilicity of the 70:30 and 50:50 PCL:SIS blended wraps increases the longevity of the release, **Figure 5C**.

**Figure 5:**
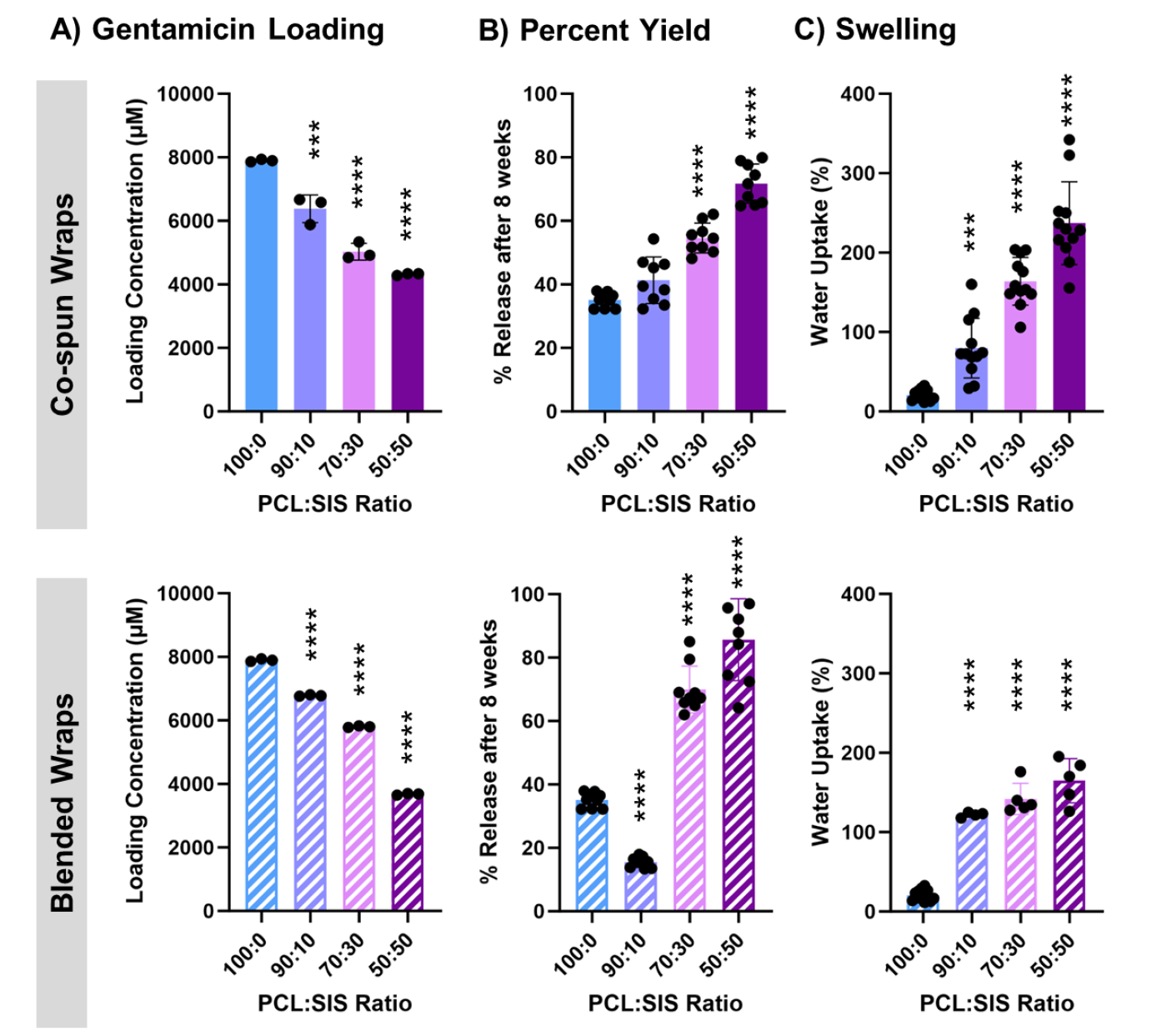
The impact of SIS incorporation on gentamicin release from PCL fibers in co-spun and blended wraps. A) Gentamicin loading concentration. B) Percent of gentamicin released from wraps after 8 weeks. C) Swelling percentage.

**Figure 6:**
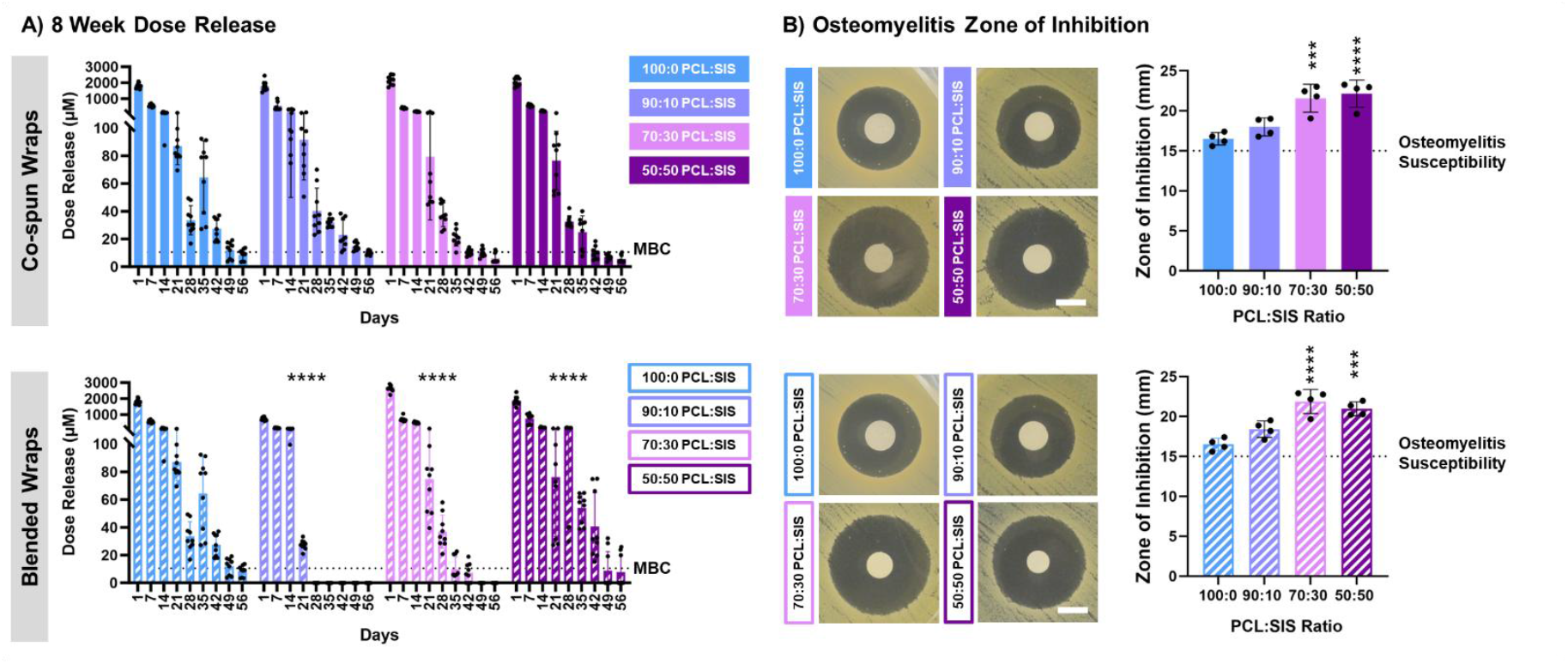
The impact of SIS incorporation on antimicrobial activity. A) Weekly release of gentamicin. The dotted line represents the minimum bactericidal concentration (MBC). B) Zone of inhibition created in osteomyelitis-specific MSSA.

A Kirby Bauer disc diffusion assay was also conducted to evaluate the impact of wrap composition on antimicrobial activity. The wraps were placed directly on an agar plate with osteomyelitis-specific *Staphylococcus aureus* (ATCC, strain 49230). The co-spun and blended wraps exhibited an increased zone of inhibition with increasing SIS incorporation, corresponding to the burst release of gentamicin, **Figure 6B**. Each wrap demonstrated a zone of inhibition > 15 mm in diameter, which indicates that *Staphylococcus aureus* is susceptible to the released gentamicin, according to the Performance Standard for Antimicrobial Susceptibility Testing.^19^ The Kirby Bauer disc diffusion assay was also conducted with wraps that had been incubated in water at 37°C for up to 8 weeks to evaluate the retention of antimicrobial activity over time, **Figure S6**. The zone of inhibition formed by the co-spun wraps remains elevated as compared to the PCL wrap and increased with increasing SIS incorporation out to 4 weeks. Still, the zone of inhibition for all co-spun wraps was reduced below that of PCL during the 8-week timepoint. Similarly, the blended wraps maintained an elevated zone of inhibition compared to the PCL wrap for 4 weeks and fell below the PCL wrap after 8 weeks. Collectively, these results indicate that SIS incorporation does not significantly impede gentamicin release from the co-spun wraps, unlike the 90:10 and 70:30 blended wraps that do not sustain release. Instead, co-electrospinning PCL and SIS improves the efficiency of release and meets infection control design criteria for up to 6 weeks.

### 3.4 Effect of PCL Incorporation on Angiogenic Properties

The primary motivation to integrate SIS into the PCL wrap was to impart angiogenic properties. After determining that the SIS incorporation did not sacrifice the durability and antimicrobial properties of the PCL wrap, the angiogenic properties of the composite wraps was compared to the pro-angiogenic SIS wrap control. *In ovo* vascularization in response to the wraps was assessed through a CAM assay, **Figure 7**. Only the 50:50 blended wrap was evaluated in the *in ovo* model to ensure animal welfare and resource conservation. The PCL wrap control resulted in a small increase in the number of intersecting vessels after 4 days. This was attributed to a modest inflammatory response associated with PCL and placing a foreign body on the CAM surface. The co-spun wraps exhibited increased angiogenic properties with increasing SIS incorporation, **Figure 7A**. The 90:10 PCL:SIS co-spun wrap resulted in a similar vascular response to the PCL wrap, indicating that 10% SIS incorporation was not sufficient to impart significant angiogenic properties. The 70:30 and 50:50 PCL:SIS co-spun wraps resulted in increased angiogenesis after 4 days. The 70:30 and 50:50 PCL:SIS wraps exhibited 36 ± 13% and 51 ± 23% of the change in intersecting vessels of the SIS wrap control, respectively, corresponding to the relative SIS mass fraction in the sample. This indicates that the angiogenic response was primarily dictated by the mass fraction of SIS present in the wrap and not negatively impacted by PCL incorporation. Interestingly, the 50:50 PCL:SIS blended wrap did not initiate vessel formation in the CAM assay, **Figure 7B**. Before placing the samples on the CAM, the samples must be hydrated to prevent damaging the CAM by wicking away water. In the 15 min of swelling prior to CAM placement, it is expected that the majority of the SIS was released from the blended wrap and was not present during the assay. The blended 50:50 wrap lost 62 ± 4% mass after swelling for 1 hour, and FTIR analysis confirmed a significant reduction in SIS after swelling, **Figure S7**. Fast resorption of digested SIS is a commonly reported challenge; however, SIS digestion is required for blend electrospinning with over 10% ECM.^10, 23^ Although the blended wrap could be implanted in the MIMT dry, these results indicate that its biological impact may be short-lived and largely guided by a burst release of SIS in the first hour after implantation. Overall, the co-spun wraps displayed increased angiogenic properties with increasing SIS incorporation and outperformed the blended wrap.

**Figure 7:**
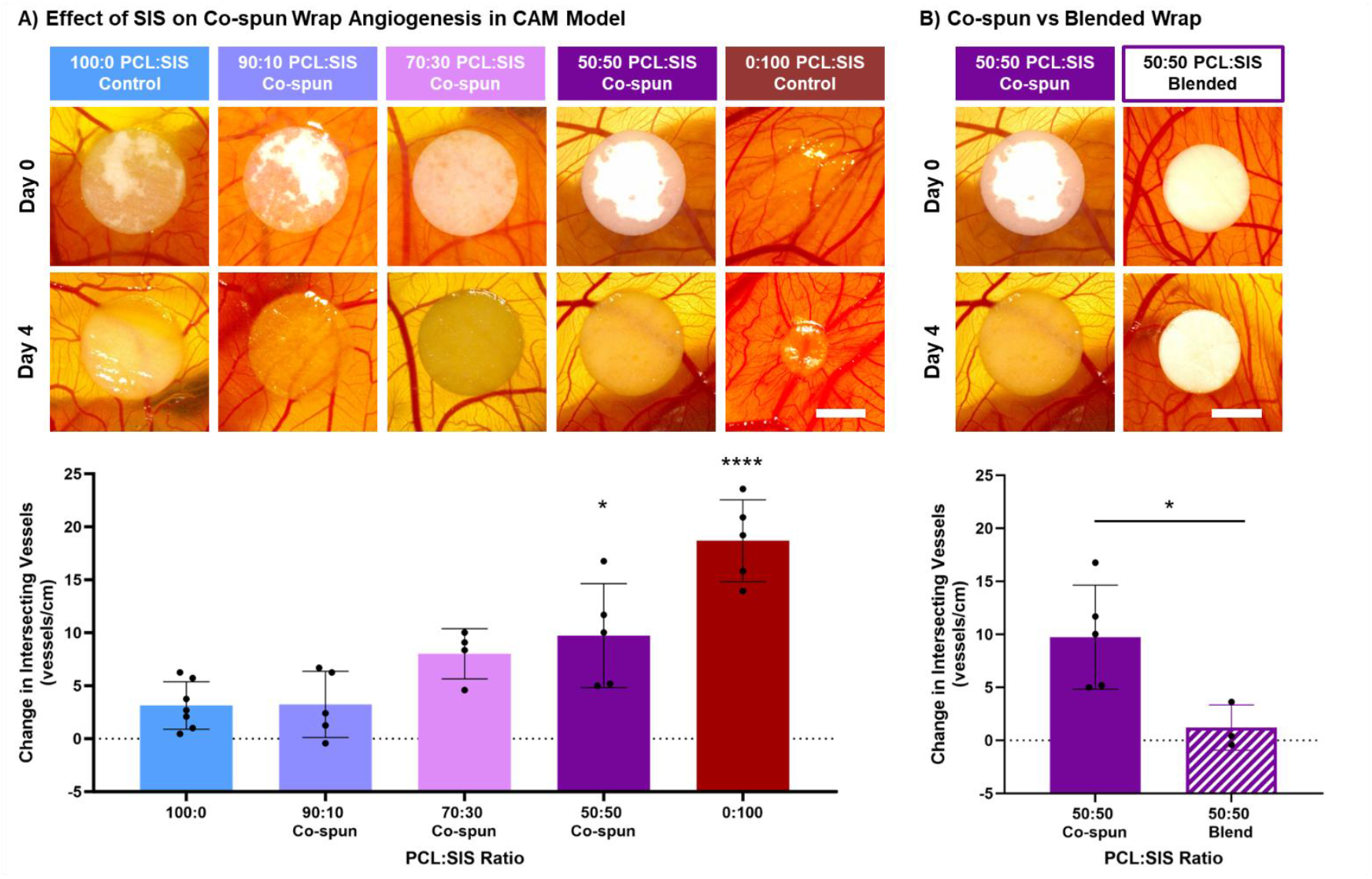
The impact of PCL incorporation on SIS angiogenic properties. A) *In ovo* angiogenesis of co- spun wraps in a CAM model after 4 days of sample placement. Scale bar = 3 mm. Vessel density is calculated as the change in the number of vessels intersecting with the sample from Day 0 to Day 4, normalized to the sample perimeter. B) Comparison between the angiogenic properties of co-spun and blended wraps.

### 3.5 Study limitations and future work

These studies describe a direct comparison of co-electrospinning versus blend electrospinning of a multifunctional PCL:SIS wrap for use in MIMT. The recent development of a suspension electrospinning method to create SIS fibers without a polymer carrier facilitated a robust characterization of the impact of SIS localization within an electrospun wrap.^17^ To our knowledge, this is the first report of a co-spun PCL:SIS wrap without the use of a polymer carrier to aid SIS spinning. Co-electrospinning and blend electrospinning were used to fabricate wraps with paired compositional ratios of PCL and SIS, measured by mass ratio and chemical composition. Due to the limitations of blend electrospinning, the level of SIS processing and fiber diameter of the wraps were not matched between the co-spun and blended wraps. Pepsin digestion was required to solubilize the SIS and load greater than 10% SIS into the blended wraps, which can lead to denaturation and impact SIS bioactivity.

Additionally, the increased solution conductivity instilled by the SIS incorporation imparted jet instability and prevented uniform collection of fibers greater than 1.5 µm in fiber diameter. Thus, the fiber diameter of the blended wraps in this study was paired to the SIS fibers within the co-electrospun wrap instead of the PCL fibers. The altered SIS processing and fiber diameter is a limitation of this study but also highlights the constraints of the blend electrospinning setup.

This work introduced advanced fabrication techniques that enable independent and modular control of competing design requirements. The resulting multifunctional wrap has demonstrated effectiveness *in vitro* by supporting surgical handling, eliminating bacteria, and enhancing angiogenesis. However, further *in vivo* orthopedic assessment and continued material refinement are necessary prior to clinical implementation. The multifunctional wrap was designed to improve the handling properties of the induced membrane. We have shown that the co-spun wrap can withstand suturing while supporting cell infiltration. One challenge of this work was establishing success criteria for surgical handling. The ISO 7198 suture retention standard was helpful, but it does not fully encompass the properties needed for a material to be considered “easy to handle” by a surgeon. SIS incorporation was shown to incrementally reduce elongation properties and increase stiffness in the PCL:SIS co-electrospun wraps, so it is essential to establish more precise criteria for the minimum flexibility needed to support surgical handling and bone wrapping. Future studies that interview surgeons about their surgical technique and investigate the forces applied during surgery would be beneficial to determine more precise handling criteria.

A primary focus of this work was also on establishing a method to prolong the release of antibiotics long-term to reduce infection recurrence and persistence. Although the potential of the PCL:SIS wrap to release gentamicin for up to 6 weeks was demonstrated, further work is needed to validate this approach and integrate it with clinical practice. Current studies were limited to *in vitro* assessment of release kinetics and bacterial susceptibility. Future studies are needed to evaluate the efficacy of gentamicin-loaded wraps to treat infection in an *in vivo* contaminated osseous defect model.

In addition, it is common for surgeons to select multiple antibiotics for local treatment, including gentamicin, vancomycin, and tobramycin. The combination of these antibiotics combats both gram-positive and gram-negative bacteria strains. In the future, multiple antibiotics could be loaded into the PCL:SIS wrap for a broad infection treatment regime. Although continuing with the current antibiotics used in clinical practice is the first step for translation of the multifunctional wrap, antibiotic resistance is a rising concern. Our lab previously determined that gallium maltolate was an effective antimicrobial for chronic wound-specific MRSA, but it was not effective for osteomyelitis. Future work should also focus on the identification of an antimicrobial agent capable of treating osteomyelitis that does not share the risk of antibiotic resistance. A few areas of ongoing research in antimicrobials for osteomyelitis are metal ions, antimicrobial peptides, peptidomimetics, and phages.^32^

Lastly, the multifunctional wrap was designed to enhance the vascularization of the induced membrane. Although we have shown the potential of the co-spun PCL:SIS wrap to promote vessel formation, future work is needed to optimize the angiogenic properties of the multifunctional bone wrap. Current studies are underway to evaluate the efficacy of the co-electrospun PCL:SIS wrap to promote vascularization in an *in vivo* subcutaneous model. SIS was chosen as a preliminary tissue source for ECM because it is rich in growth factors that promote vascularization and is widely studied in tissue regeneration applications. However, the newly developed method to electrospin ECM from tissue sources with non-flat structures opens up the possibility of other tissue-sourced ECM. Future studies are necessary to evaluate the angiogenic properties of ECM derived from different tissue sources such as bone, vascular, and amnion tissue and identify the ECM composition that can further optimize the formation of mature and lasting vasculature in the induced membrane. In addition to optimizing ECM tissue source, the ECM suspension electrospinning method used in this work have room for improvement. Most notable is the need to transition from a harsh solvent (HFIP) to a less toxic electrospinning solvent. There is promising work on electrospinning natural materials with 20X PBS and ethanol or acetic acid that could be modified for use in the ECM suspension electrospinning setup. Once a robust method is developed to enhance the vascularity of the induced membrane, a mechanistic study is needed to evaluate the impact of vascularization on bone healing in MIMT. The mechanism of bone healing by the induced membrane remains largely unknown. Optimizing our method to modify the vessel formation, immune cell response, and membrane composition will pave the way for fundamental studies on bone healing and elucidate critical parameters for the development of a pseudo-periosteum as an adjunct therapy to bone grafts.

## 4. Conclusion

In this study, we designed an electrospun wrap to augment membrane durability, sustain infection control, and enhance membrane vascularity in MIMT. A range of PCL:SIS compositions was evaluated to balance the needs for surgical handling, antibiotic release, and angiogenic properties. Co-spun wraps demonstrated improved handling properties as compared to ECM wraps and solvent welding was used to achieve target suture retention standards without diminishing SIS content. Interestingly, the hydrophilicity imparted by SIS incorporation overcame the loss of PCL mass in the co-spun wraps and resulted in sustained gentamicin release above the MBC for up to 6 weeks, 3 times as long as the clinical standard and at a higher yield. Although the blended wraps showed similar trends to the co- spun wraps for the first two weeks, gentamicin release was not maintained out to 6 weeks. Lastly, angiogenic properties were instilled into the co-spun wraps with at least 30% SIS mass fraction, and the *in ovo* vascular response corresponded to the percent mass of SIS in the co-spun wraps. The blended wrap did not trigger an angiogenic response due to the premature burst release of SIS. These results highlight the advantage of a co-spun system over a blended system. The modular nature of co-electrospinning facilitates greater control over fiber diameter, inter-fiber fusion, and SIS digestion not afforded by a blended system. Altogether, the 50:50 co-spun PCL:SIS wrap met the design criteria for surgical handling and antibiotic release while exhibiting the maximum angiogenic response. Future work will evaluate this bone wrap in an *in vivo* MIMT model to characterize the handling, antimicrobial, and angiogenic properties of the wrap and the subsequent impact on bone healing. Overall, this study established the benefit of a co-electrospun modular design and the efficacy of using a co-spun fiber PCL:SIS wrap to simultaneously augment membrane durability, sustain infection control, and enhance membrane vascularity.

## 6. Acknowledgements

Funding for this work was supported by the National Science Foundation Graduate Research Fellowship Program under Grant (award # 2020300397). Any opinions, findings, conclusions, or recommendations expressed in this material are those of the author(s) and do not necessarily reflect the views of the National Science Foundation.

