## Supplemental Information for "Co-Electrospinning Extracellular Matrix with Polycaprolactone Enables a Modular Approach to Balance Bioactivity and Mechanics of a Multifunctional Bone Wrap"

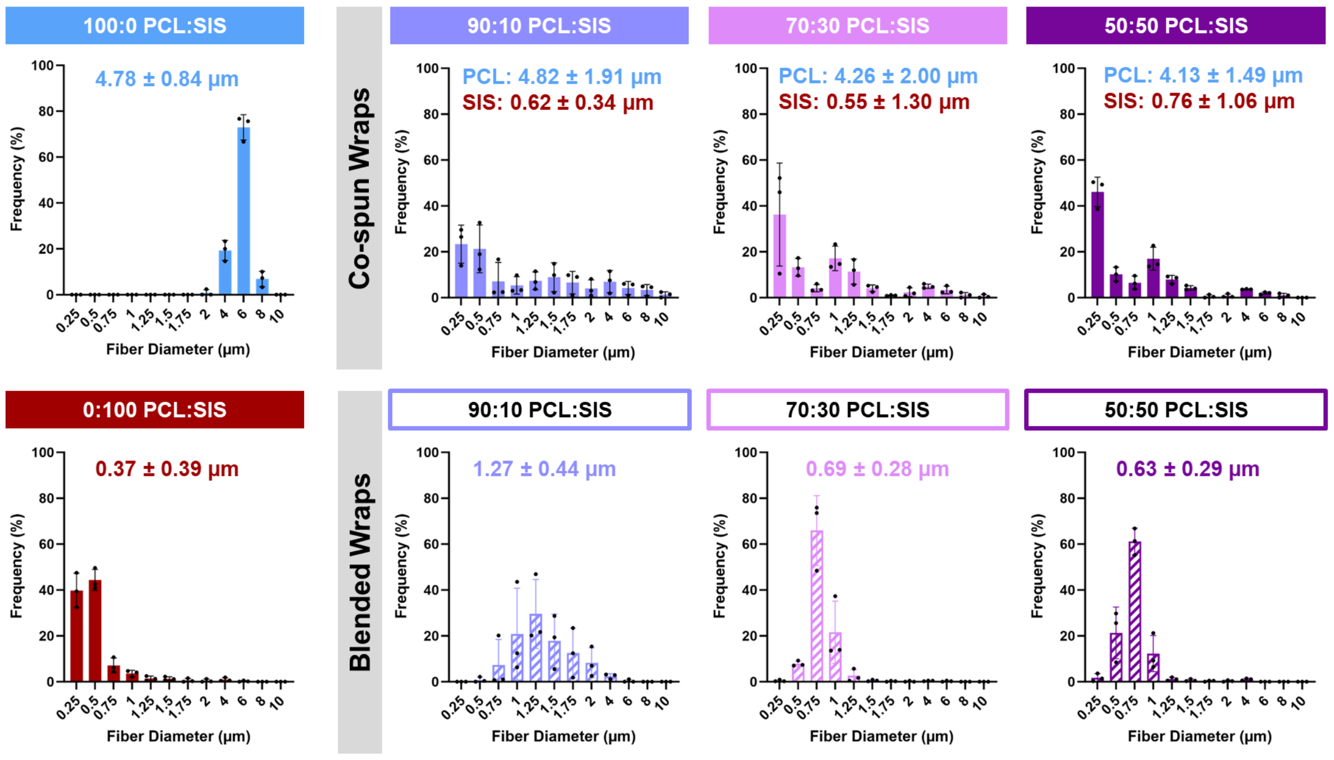


***Figure S1:*** Fiber diameter analysis of co-spun and blended wraps. The fiber diameter is reported as average ± standard deviation. The fiber diameters of the dual fiber wraps are calculated separately for each population, categorized as either less than or greater than 2 µm, corresponding to the SIS and PCL fibers, respectively.

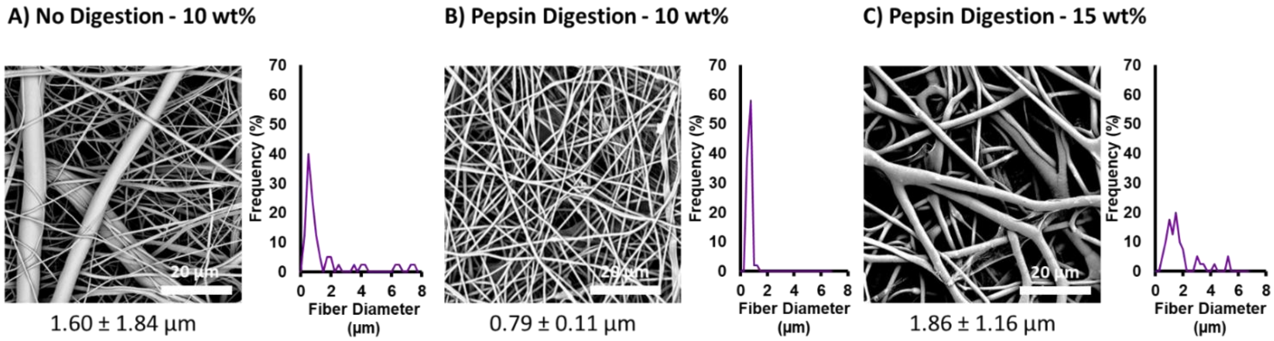


***Figure S2:*** Scouting electrospinning parameters for blended fiber wraps to increase fiber diameter and reduce fiber splitting. A) 10 wt% PCL:SIS solution concentration without SIS digestion. B) 10 wt% PCL:SIS solution concentration with 72 h pepsin digestion. C) 15 wt% PCL:SIS solution concentration with 72 h pepsin digestion. Scanning electron micrographs (left) and a frequency distribution of fiber diameter (right) of each wrap. The fiber diameter is listed as average ± standard deviation.

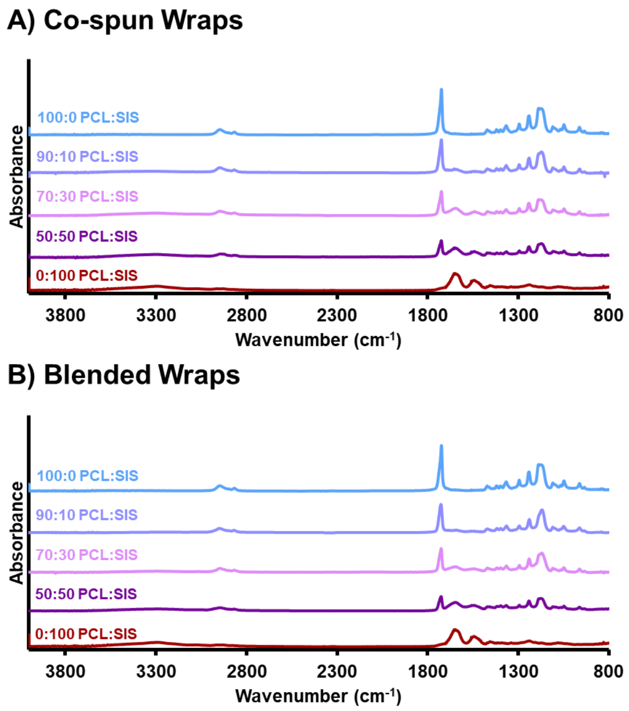


***Figure S3:*** Full ATR-FTIR spectra of co-spun (A) and blended (B) PCL:SIS wraps.

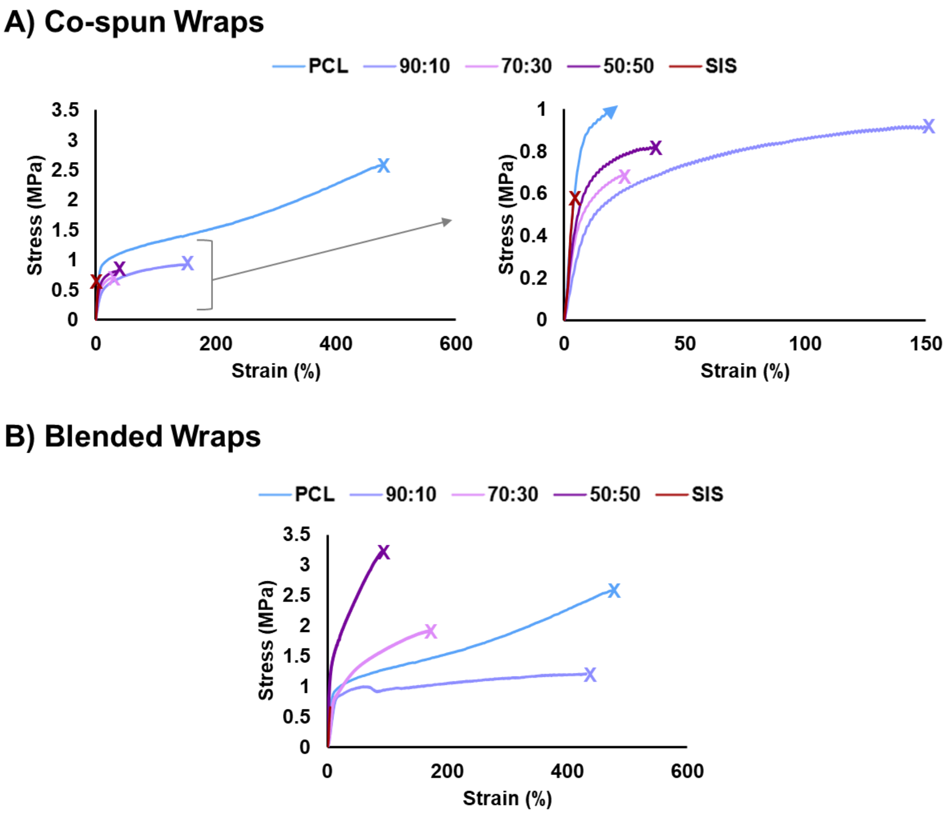


***Figure S4:*** Uniaxial tensile stress-strain curves of co-spun (A) and blended (B) PCL:SIS wraps.

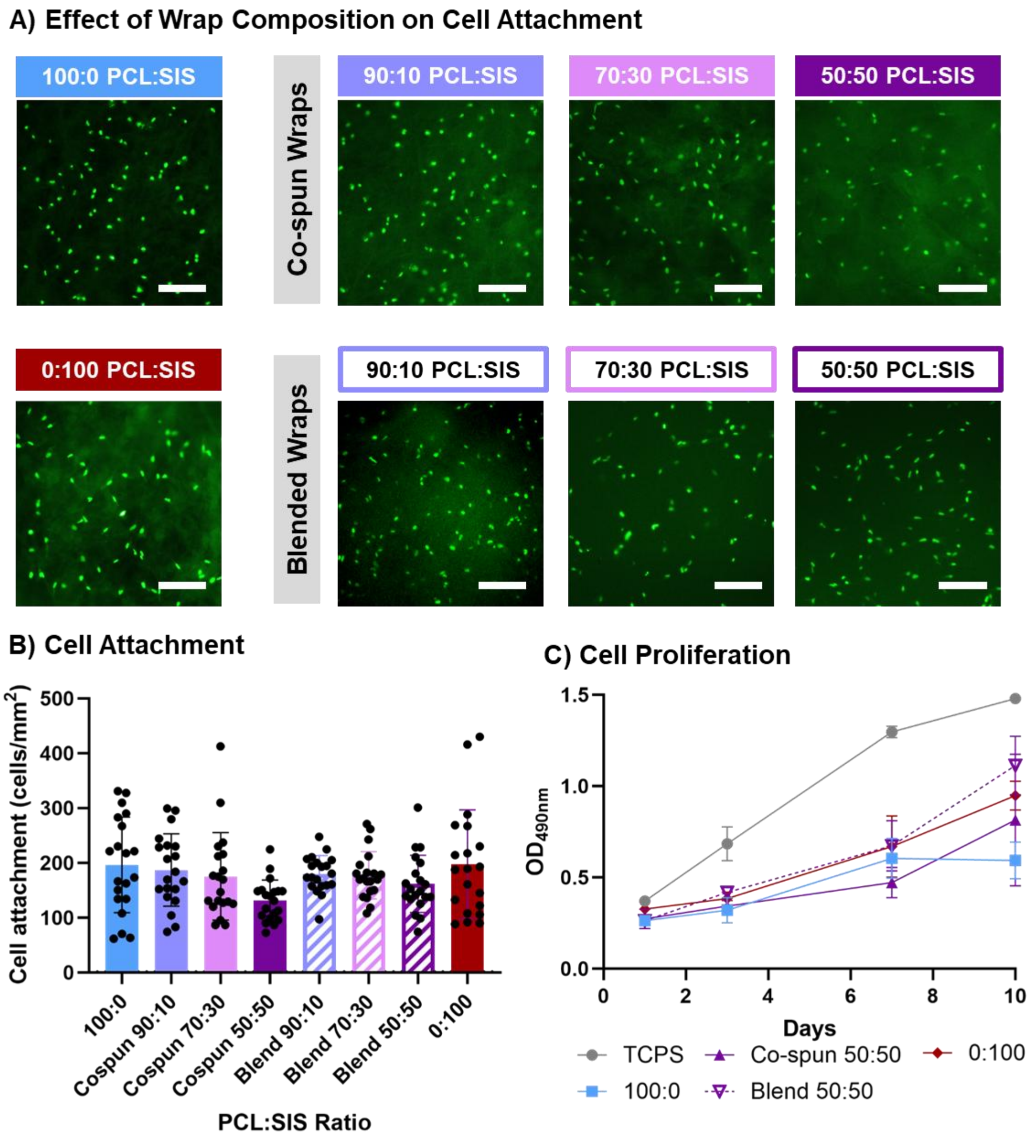


***Figure S5:*** Human dermal fibroblast attachment, viability, and proliferation on co-spun and blended PCL:SIS wraps. A-B) Cell attachment on the surface of the wraps after 3 hours. Scale bar = 300 µm. C) Cell proliferation as assessed by metabolic activity after 1, 3, 7, and 10 days of culture.

To further validate the sustained antimicrobial activity of the composite wraps, the zone of inhibition assay was adapted to assess wraps after incubation in water over 8 weeks **(Figure S6)**. The composite wraps were cut into 6 mm discs and incubated in 2 mL of DI water for 1, 4, and 8 weeks with weekly water changes. At the end of each timepoint, the specimens were washed in 2 mL of DI water and dried on the benchtop overnight. The antimicrobial capacity of the incubated specimens was then evaluated with the zone of inhibition assay by placing the meshes on the bacterial plate.

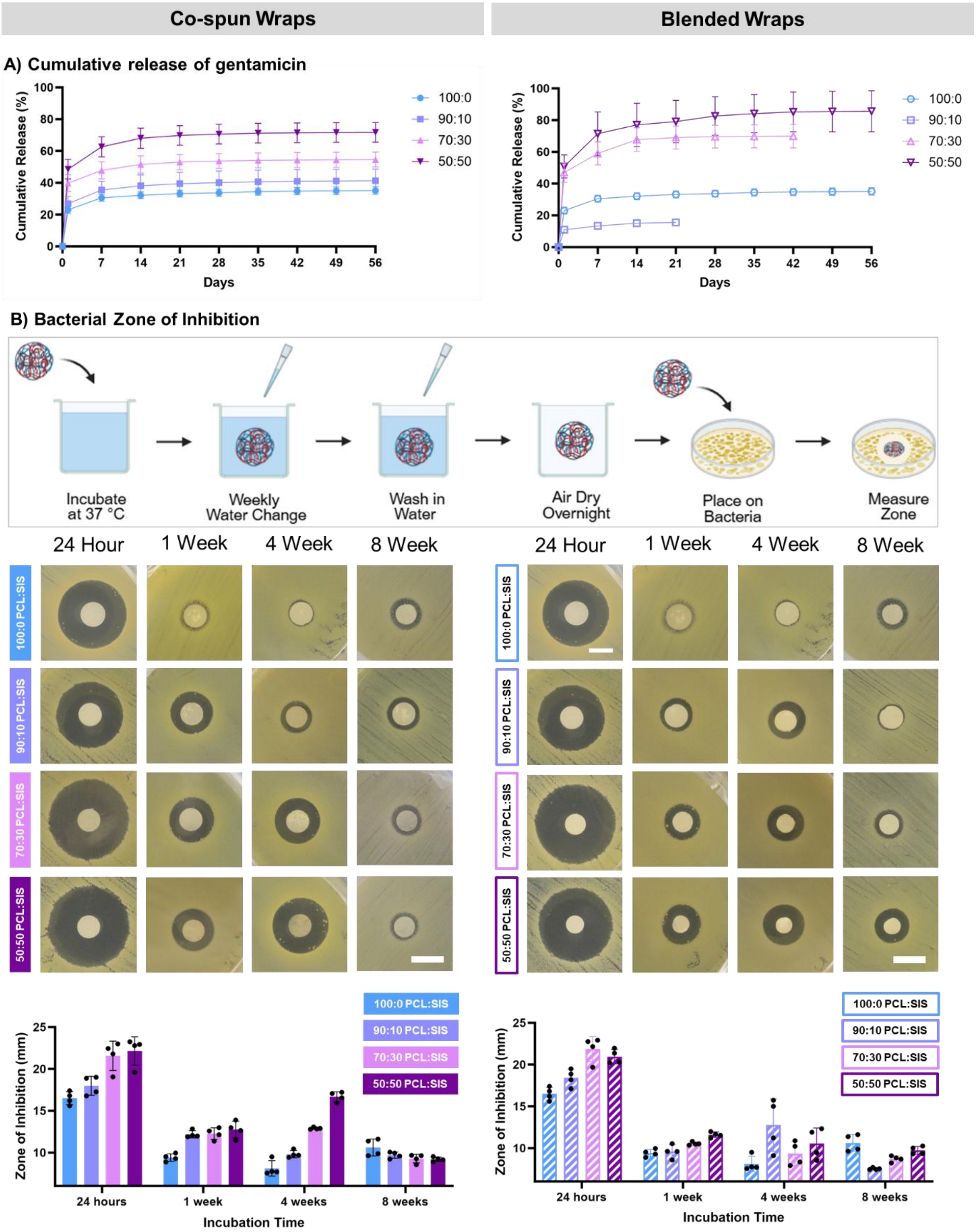


***Figure S6:*** Antimicrobial properties of co-spun and blended wraps. A) Cumulative release of gentamicin from wraps over 8 weeks. B) Kirby Bauer disc diffusion assay of co-spun and blended PCL:SIS wraps after 0 – 8 weeks of incubation in water. Created in BioRender. Jones, S. (2025) https://BioRender.com/m2697ke

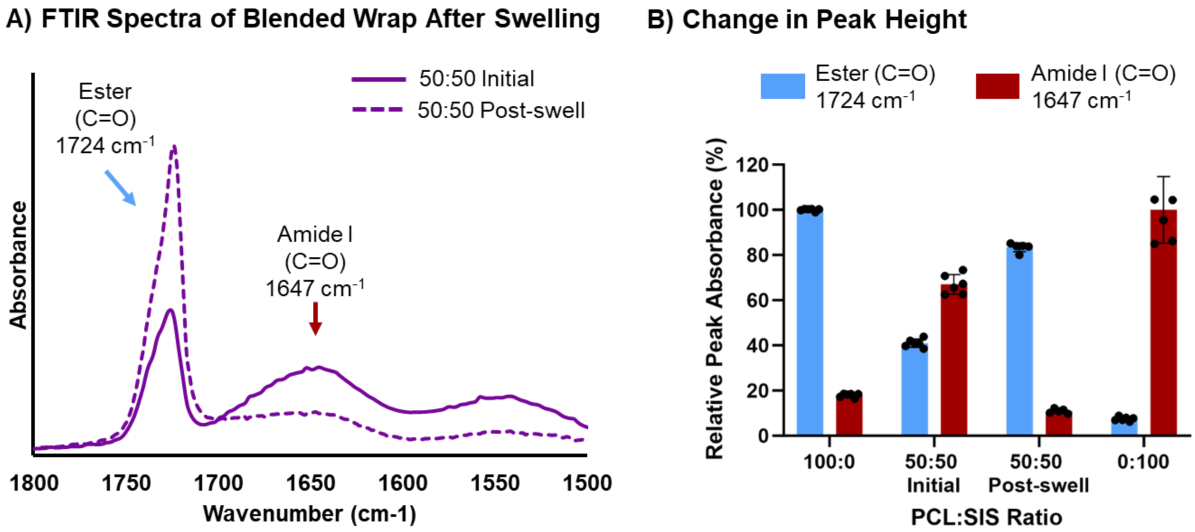


***Figure S7:*** ATR-FTIR spectral analysis of the 50:50 PCL:SIS blended fiber wrap before and after swelling in water for 1 hour. A) ATR-FTIR spectra with B) relative peak height absorbance of ester (C=O, 1724 cm$^{-1}$) and amide I (C=O, 1647 cm$^{-1}$) normalized to electrospun PCL and SIS controls.
